# Msp1-dependent extraction promotes ubiquitylation of translocation-stalled mitochondrial precursor proteins

**DOI:** 10.64898/2026.09.28.754956

**Authors:** Jeannine Engelke, Noemi Occhipinti, Lisanne Pitzke, Kira Ritzenhofen, Robert Hardt, Marc Sylvester, Enzo Scifo, Fabian den Brave, Thomas Becker

**Affiliations:** Institute of Biochemistry and Molecular Biology, University Hospital Bonn, University of Bonn, 53115 Bonn, Germany; Core Facility Analytical Proteomics, University Hospital Bonn, University of Bonn, 53115 Bonn, Germany

**Keywords:** Protein import, mitochondria, outer membrane, protein degradation, mitoTAD, mitoCPR, TOM complex, ubiquitin proteasome system

## Abstract

The translocase of the outer membrane (TOM complex) imports more than 1,000 proteins into mitochondria. Clogging of the TOM pore with a precursor protein causes proteotoxic stress and eventually cell death. Two quality control pathways remove translocation-stalled precursor proteins. In the mitochondrial protein translocation-associated degradation (mitoTAD), Ubx2 recruits the cytosolic AAA-ATPase Cdc48 to clear precursor proteins from the TOM complex. In the mitochondrial compromised protein import response (mitoCPR), the stress-induced Cis1 recruits the AAA-ATPase Msp1 to Tom70. The role of Msp1 for the removal of mitochondrial precursor proteins remains unknown. Here, we demonstrate that parallel loss of Msp1 and Ubx2 strongly affects removal of precursor proteins and cell viability. Msp1 and Ubx2 bind independently of import stress and Cis1 to the TOM complex to remove a large variety of precursor proteins. Msp1-dependent extraction promotes ubiquitylation of precursor proteins, which in turn allows Ubx2-recruited Cdc48 to transfer the substrates to proteasomal degradation. We conclude that two AAA-ATPases cooperate in mitochondrial precursor quality control. Msp1-dependent extraction from the TOM complex facilitates precursor ubiquitylation and Cdc48-mediated transfer to proteasomal degradation.

## INTRODUCTION

Mitochondria import about 1,000 proteins in bakeŕs yeast *Saccharomyces cerevisiae* and 1,500 proteins in humans^1,2^. The proteins are made as precursors on cytosolic ribosomes and are guided by a network of molecular chaperones to the mitochondrial surface^3,4^. The translocase of the outer membrane (TOM complex) imports almost all precursor proteins into the organelle. After passage of the TOM pore, specific protein translocases sort the precursor proteins into the mitochondrial subcompartments: matrix, intermembrane space, inner and outer membranes^5–8^. About 60% of the mitochondrial proteins are produced with a cleavable presequence that is removed upon import^5,7,9^. The presequence translocase (TIM23 complex) transports these proteins into the matrix or inserts them into the inner membrane. The mitochondrial processing peptidase (MPP) cleaves the presequence of imported proteins to allow their folding. All other mitochondrial proteins contain non-cleavable sorting signals that direct them to their respective mitochondrial subcompartments^5–8^.

Protein import into mitochondria is a challenge for cellular homeostasis^10–17^. Impaired protein import into mitochondria causes accumulation of precursor proteins, leading to proteotoxic stress and eventually cell death^18–23^. Non-imported mitochondrial precursor proteins can be sorted into protein deposits, termed MitoStores, transported to different cellular compartments and are eventually degraded by the proteasome^24–28^. The accumulation of non-imported precursor proteins induces a transcriptional reprogramming, leading to increased expression of genes encoding for chaperones and quality control components to counteract the proteotoxic stress. In contrast, the expression of genes encoding for components involved in oxidative phosphorylation is decreased to reduce the load of proteins that have to be imported into mitochondria^18–20^.

Clogging of the TOM channel with a precursor protein impairs protein import into mitochondria^20,24,29^. Precursors can arrest in the TOM channel during translocation for different reasons. First, a substantial fraction of the newly synthesized proteins fails to adopt their mature conformation^30^. Misfolding or premature folding of precursor proteins can prevent their translocation into mitochondria since largely folded domains cannot pass the TOM pore^31^. Second, polypeptide chains that are not released from the ribosome can accumulate in the translocation pore^32^. Third, the activity of the respiratory chain generates a membrane potential across the inner membrane that drives protein translocation via the TIM23 complex^33,34^. Mitochondrial dysfunction can lead to a depletion of the membrane potential, which in turn blocks the import of precursor proteins across or into the inner membrane. Finally, defects of the translocon can also delay protein transport^19,24^. Therefore, mitochondrial biogenesis and function depend on mechanisms that clear translocation-arrested precursor proteins from the TOM channel^10,11,13–16^. Two quality control pathways at the TOM complex have been identified^24,29^. The mitochondrial protein translocation-associated degradation (mitoTAD) pathway constitutively monitors protein import^24^. The core subunit Ubx2 associates with the TOM complex and recruits the cytosolic AAA-ATPase Cdc48. The hexameric Cdc48 powers the extraction of precursor proteins for subsequent proteasomal degradation^24^. Similarly, Ubx2 constitutes a docking site for Cdc48 in the endoplasmic reticulum in the ER-associated degradation pathway (ERAD)^35,36^. The mitochondrial compromised protein import response (mitoCPR) comprises the import stress-triggered expression of the cytosolic protein Cis1. Overproduced Cis1 recruits the outer membrane-bound AAA-ATPase Msp1 to Tom70 to extract precursor proteins from the TOM complex^28,29^. The extracted protein can either be degraded or imported^29,37^. According to the current view, mitoTAD and mitoCPR act independently of each other. How the activities of both AAA-ATPases, Ubx2-recruited Cdc48 and Msp1, are coordinated under different physiological conditions remains unclear.

Non-imported mitochondrial precursor proteins are degraded by the ubiquitin-proteasome system. The ubiquitin-proteasome system involves polyubiquitylation of client proteins by an enzymatic cascade to label them for proteasomal degradation^38^. Ubiquitin is a highly conserved 76 amino acid long polypeptide. E3 ubiquitin ligases mediate the transfer of activated ubiquitin onto the client proteins. The E3 ubiquitin ligases Rsp5 in yeast and MARCH5 in mammals, as well as the deubiquitylase Ubp16 and its mammalian counterpart USP30, control ubiquitylation of non-imported precursor proteins at the TOM complex^39–41^. Pth2 constitutes a docking site for the ubiquilin Dsk2 and therefore facilitates the transfer of ubiquitylated proteins to the proteasome^41^. The role of precursor protein ubiquitylation in the mitoTAD and mitoCPR pathways is largely unknown.

We investigated a possible interplay between Ubx2-Cdc48 of the mitoTAD pathway and Msp1 of the mitoCPR pathway in the removal of precursor proteins. We found that the parallel loss of Msp1 and Ubx2 compromises the removal of non-imported mitochondrial precursor proteins, affecting cell viability. We showed that Msp1 and Ubx2 associate with the TOM complex independently of stress and Cis1. Msp1 and Ubx2 share binding to a variety of precursor proteins. Finally, we uncovered that several hundred mitochondrial precursor proteins can be ubiquitylated upon import failure, revealing the pivotal role of the ubiquitin-proteosome system in the quality control of mitochondrial protein import. Msp1 promotes ubiquitylation of these precursor proteins, while Ubx2-Cdc48 complete their extraction from the TOM complex to transfer them for degradation. Thus, Msp1 and Ubx2 do not function in separate quality control pathways, but rather display sequential roles in the degradation of non-imported precursor proteins by the ubiquitin-proteasome system. We conclude that the two AAA-ATPases, Msp1 and the Ubx2-recruited Cdc48, monitor protein transport at the TOM complex to ensure full import competence.

## RESULTS

### Msp1 and Ubx2 functionally interact in the removal of mitochondrial precursor proteins

Msp1 of the mitoCPR pathway was reported to clear precursor proteins from the TOM complex under import stress conditions^29^. Whether it cooperates with other quality control factors at the TOM complex remains unknown. To study a possible functional interplay, we generated double deletion strains of *msp1Δ* combined with either *ubx2Δ*, *vms1Δ* or *pth2Δ* (Fig. 1a). Vms1 is a peptidyl-tRNA hydrolase that functions in the ribosome-associated quality control of mitochondrial proteins^32,42^. Excitingly, only the *msp1Δ ubx2Δ* strain displays a strong growth defect, while the other double deletions did not show any synthetic growth defect (Fig. 1b)^24^. We wondered whether the parallel loss of Msp1 and Ubx2 impairs the degradation of precursor proteins. If the import is impaired, mitochondrial precursor proteins with a cleavable presequence are not processed by MPP, resulting in the detection of the larger precursor form in addition to the imported mature form. Analysing total cell extracts, we detected an accumulation of the matrix targeted precursor proteins of Ilv2 and Mdj1 in *msp1Δ ubx2Δ* cells (Fig. 1c). To investigate whether the degradation of these proteins is impaired, we treated the cells with the ionophore carbonyl cyanide m-chlorophenyl hydrazine (CCCP) to deplete the mitochondrial membrane potential and therefore block protein import. Under these conditions, non-imported precursor forms of Ilv2 and Atp2 accumulate compared to untreated cells (Fig. 1d). Subsequently, the cells were treated with cycloheximide to block the de novo protein synthesis. The Ilv2 and Atp2 precursors are degraded over time in wild-type cells (Fig. 1d). While the degradation of these precursor proteins was only mildly delayed in the *msp1Δ* and *ubx2*Δ cells, the proteins were strongly stabilized in the double deletion cells (Fig. 1d). We next investigated whether clogging of the mitochondrial translocon affects cell growth. Therefore, we overexpressed cytochrome *b_2_*(1-84)-fused to DHFR followed by the heme binding domain of cytochrome *b*_2_ (*b*_2_-DHFR) and cytochrome *b*_2_(1-84)Δ-DHFR that lacks the hydrophobic inner membrane sorting signal (amino acids 47-65) for insertion into the inner membrane (*b*_2_Δ-DHFR). When overexpressed both precursor proteins arrest in the TOM complex and clog the import channel^24^. Remarkably, the expression of both clogger proteins is lethal for the double deletion *msp1Δubx2*Δ, while the growths of the single deletion strains were affected to a similar extent as wild-type (Fig. 1e). The protein content of isolated *msp1Δ ubx2*Δ mitochondria, including TOM and respiratory chain complexes, remains largely unaffected (Extended Data Fig. 1), excluding the possibility that reduced amounts of the TOM complex lead to the accumulation of the precursor proteins. Thus, Ubx2 and Msp1 can partially compensate for each other, however, the parallel loss of both factors strongly impairs removal of non-imported precursor proteins.

**Figure 1.**
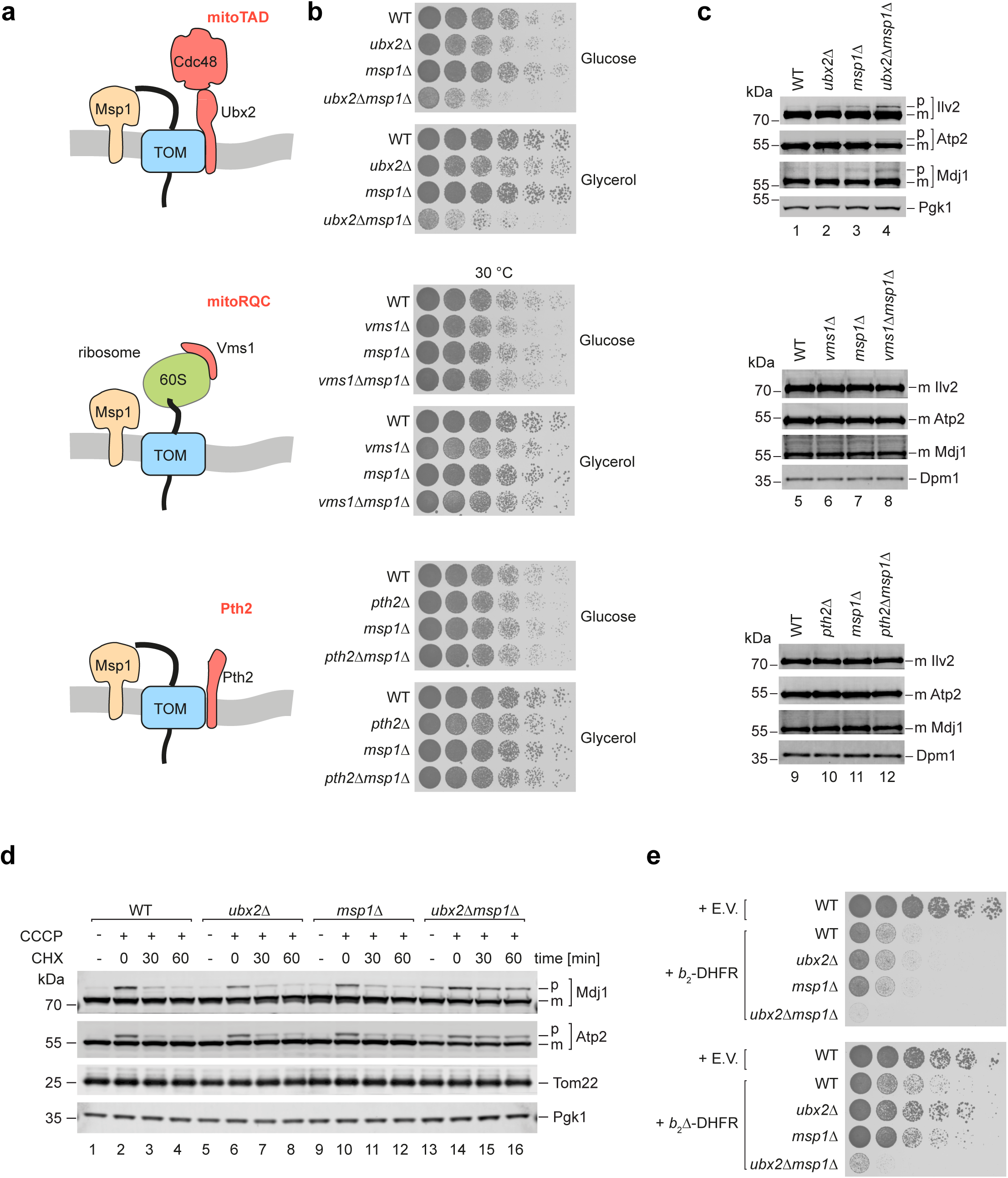
Msp1 genetically interacts with Ubx2. **(a)** Scheme depicting the different quality control factors acting on precursor proteins at the TOM complex. **(b)** Serial dilutions of wild-type (WT), *ubx2Δ, msp1Δ* and *ubx2Δ msp1Δ* strains were grown on media containing the indicated carbon source at 30°C. **(c)** Cell extracts of WT, *ubx2Δ, msp1Δ* and *ubx2Δ msp1Δ* strains were analysed by Western blotting using the indicated antisera. p: precursor; m: mature. **(d)** WT, *ubx2Δ, msp1Δ* and *ubx2Δ msp1Δ* cells were treated with CCCP for 1 h. Where indicated, cells were treated with cycloheximide (CHX) for the indicated time points. Cell extracts were analysed by immunoblotting with the indicated antisera. p: precursor; m: mature. **(e)** Serial dilutions of the indicated strains expressing cytochrome *b_2_*(1-84)-fused to DHFR followed by the heme binding domain of cytochrome *b*_2_ (*b*_2_-DHFR) or its variant lacking the hydrophobic sorting signal (*b*_2_Δ-DHFR) were grown in selective media containing galactose at 30°C. E.V., empty vector.

### Msp1 and Ubx2 bind independently of import stress to the TOM complex

The observation that the combined loss of Msp1 and Ubx2 strongly affects the quality control of non-imported mitochondrial precursor proteins was surprising, since both quality control factors have been reported to function under specific conditions: Ubx2 binds robustly to the TOM complex and recruits Cdc48 to monitor protein entry into mitochondria under constitutive conditions, while Msp1 clears the clogged TOM complex under stress conditions^24,29^. To tackle this conundrum, we analysed the binding of both factors to the TOM complex under import stress and non-stress conditions. We optimized the affinity purification of HA-tagged Tom40 from mitochondria and cellular extracts using the non-ionic detergent digitonin to maintain labile protein-protein interactions. We studied the binding of quality control factors to the TOM complex in the absence of import stress by mass spectrometry (Fig. 2a and Supplementary Table 1). To exclude any loss of protein-protein interactions during isolation of mitochondria^24^, we also performed a Tom40_HA_ pulldown from total cell extracts and analysed it via Western blotting (Fig. 2b). Under these non-stress conditions, we could detect both Msp1 and Ubx2 as binding partners of Tom40 in mitochondrial and cellular extracts. However, Msp1 was co-purified with reduced efficiency compared to Ubx2 (Figs. 2a and b) as previously reported^24,29^.

**Figure 2.**
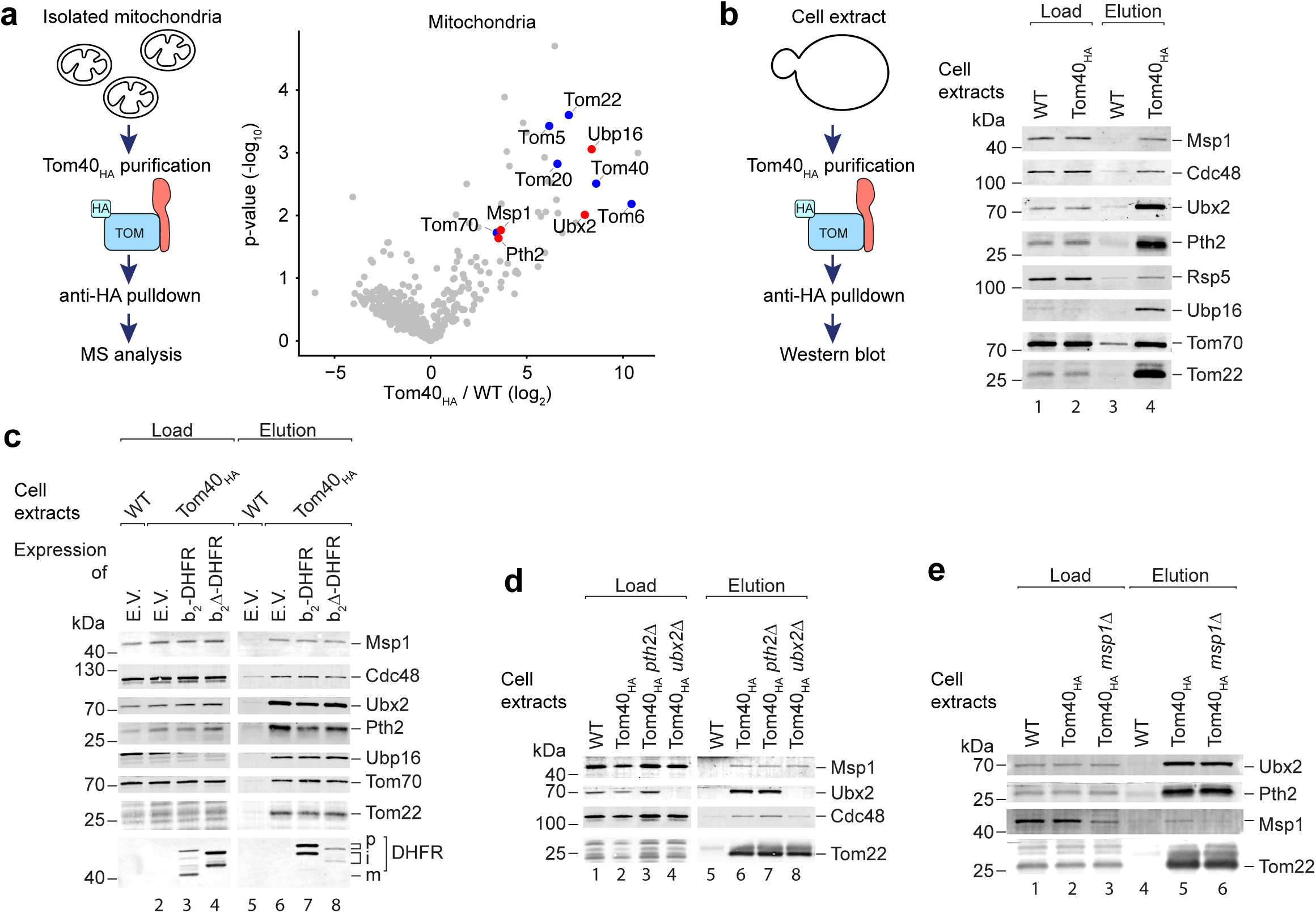
Msp1 and Ubx2 bind constitutively to the TOM complex. **(a)** Left: Scheme of experimental setup for the purification of Tom40_HA_ of the TOM complex from isolated mitochondria. Right: Mitochondria from wild-type (WT) and Tom40_HA_ cells were subjected to affinity purification. The elution fractions of three independent replicates were analysed by mass spectrometry. The log_2_ (fold change) of Tom40_HA_ versus WT is plotted against the -log10 (p-value). TOM subunits are highlighted in blue, TOM-associated quality control factors in red. **(b)** Left: Scheme of experimental setup for the purification of Tom40_HA_ from cell extracts. Right: Affinity purifications from WT or Tom40_HA_ cell extracts were analysed by immunoblotting using the indicated antisera. Load 0.2 %, elution 100 %. **(c)** Affinity purification from WT or Tom40_HA_ cells expressing cytochrome *b_2_*(1-84)-fused to DHFR followed by the heme binding domain of cytochrome *b*_2_ (*b*_2_-DHFR) or its variant lacking the hydrophobic sorting signal (*b*_2_Δ-DHFR) were analysed by immunoblotting with the indicated antisera. Load 0.2 %, elution 100 %. p: precursor; i: intermediate; m: mature. **(d)** Affinity purification from WT, Tom40_HA_, Tom40_HA_ *ubx2Δ* and Tom40_HA_ *pth2Δ* cell extracts were analysed by immunoblotting with the indicated antisera. Load 0.2 %, elution 100 %. **(e)** Affinity purification from WT, Tom40_HA_, and Tom40_HA_ *msp1Δ* cell extracts were analysed by immunoblotting with the indicated antisera. Load 0.2 %, elution 100 %.

We next tested whether the interactions of Msp1 and Ubx2 with the TOM complex are affected upon import stress. Therefore, we overexpressed *b_2_*-DHFR or *b_2_*Δ-DHFR to clog the translocon and induce import stress^20,24^. Upon import, the *b*_2_-DHFR precursor is processed twice. The mitochondrial processing peptidase (MPP) removes the presequence, while the inner membrane protease (IMP) removes the inner membrane anchor, leading to the release of *b*_2_-DHFR into the intermembrane space^43^. The precursor of *b*_2_Δ-DHFR lacks the sorting signal and is transported into the matrix, where it is processed by MPP (Fig. 2c). When we overexpressed these proteins, we could detect the precursor and intermediate forms, but not the imported mature form, in the eluate of the affinity purification of Tom40_HA_ (Fig. 2c). The co-purification of neither Msp1 nor Ubx2 was affected by the presence of a clogger (Fig. 2c), revealing that both proteins bind to the TOM complex under stress and non-stress conditions. Finally, we wondered whether the binding of Ubx2 and Msp1 to the TOM complex depended on quality control factors. Thus, we performed Tom40_HA_ pulldowns in *msp1Δ*, *ubx2Δ,* and *pth2*Δ cells. However, the co-purification of Ubx2 and Msp1 was not affected in the mutant strains (Fig. 2d), indicating that the quality control factors bind independently of each other to the TOM complex.

The expression of a clogger construct induces the expression of *CIS1* as shown by increased *CIS1* transcript level (Fig. 3a) as reported^20,29^. Overexpressed Cis1 stimulates the binding of tagged Msp1 with Tom70^29^. We asked whether Cis1 is required for the co-purification of Msp1 along with HA-tagged Tom40. We performed the affinity purification from wild-type and *cis1*Δ cell extracts to maintain labile protein-protein interactions and analysed the samples via mass spectrometry and Western blotting (Fig. 3b and Supplementary Table 2). We could not detect any impact on the association of Msp1 or Ubx2 with the TOM complex in *cis1*Δ compared to wild-type cells (Figs. 3b-d; Extended Data Fig. 2). For functional analysis, we wondered whether Cis1 like Msp1 genetically interacts with Ubx2. Therefore, we generated a double deletion strain of *CIS1* and *UBX2*. Unlike *msp1Δ ubx2Δ* (Fig. 1), we could not see any synthetic growth defect of *cis1*Δ *ubx2*Δ and could not detect any accumulation or stabilization of non-imported precursor proteins in the double deletion strain (Figs. 3e-g). We conclude that Cis1 is neither required for binding of Msp1 to the TOM complex nor for the functional interplay of Msp1 with Ubx2 in the removal of precursor proteins.

**Figure 3.**
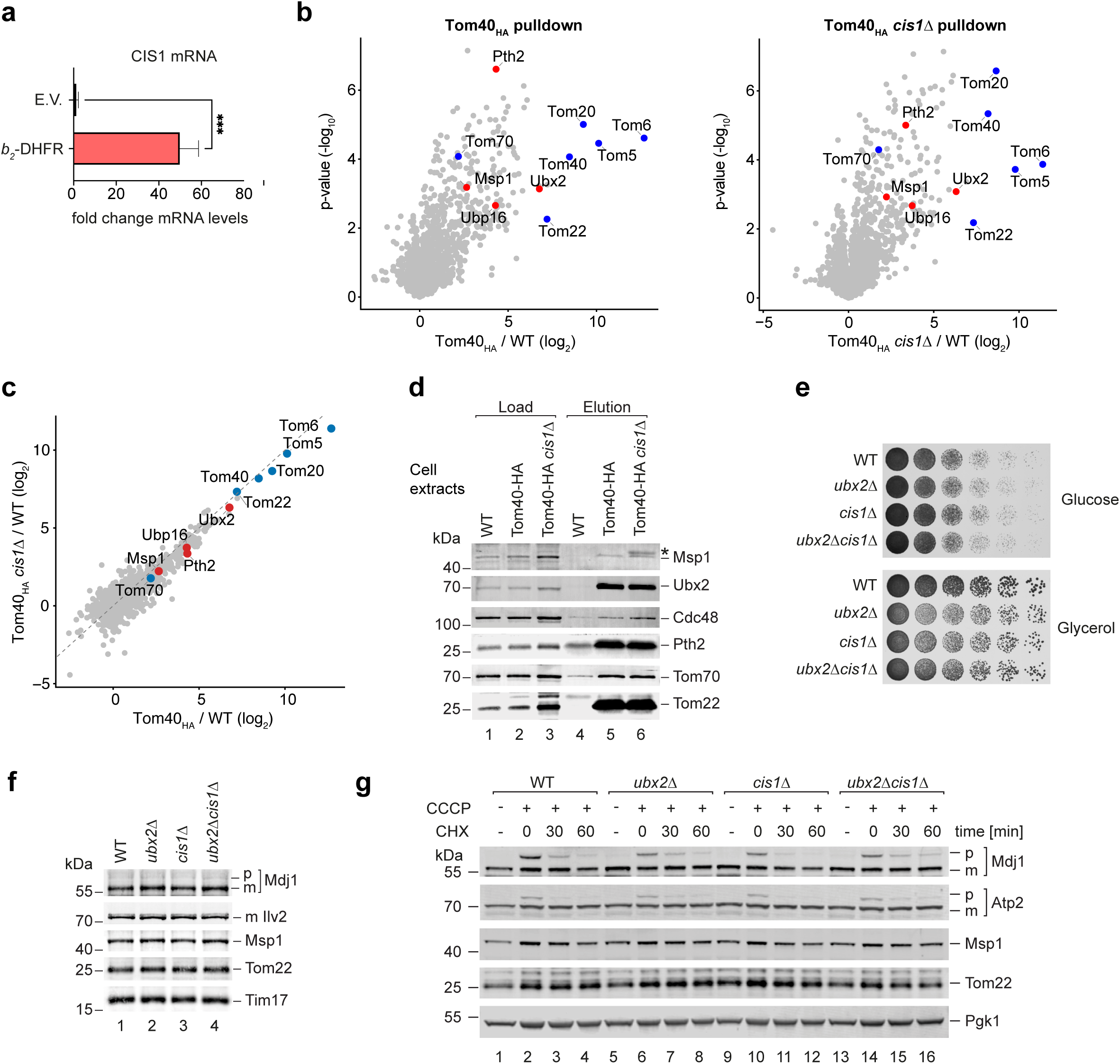
Msp1 and Ubx2 bind independently of Cis1 to the TOM complex. **(a)** Relative abundance of *CIS1* mRNA in wild-type (WT) cells with or without expression of the *b_2_*-DHFR clogger was analysed by qPCR. *CIS1* mRNA levels in wild-type with empty vector were set to 1. E.v., empty vector **(b)** Cell extracts from wild-type (WT), Tom40_HA_ and Tom40_HA_ *cis1*Δ strains were subjected to affinity purification. The elution fractions of four independent replicates were analysed by mass spectrometry. The log_2_ (fold change) of Tom40_HA_ versus WT (left) or Tom40_HA_ *cis1Δ* versus WT control (right) are plotted against the -log_10_ (p-value). Blue dots, TOM subunits; red dots, quality control factors. **(c)** The log_2_-fold change of Tom40_HA_ versus WT control (x-axis) is plotted against the log_2_-fold change of Tom40_HA_ *cis1Δ* versus WT (y-axis). Blue dots, TOM subunits; red dots, quality control factors. **(d)** Affinity purifications from WT, Tom40_HA_ and Tom40_HA_ *cis1Δ* cell extracts were analysed by immunoblotting with the indicated antisera. Load 0.2 %, elution 100 %. Asterisk denotes unspecific band of the Msp1 antiserum (see also Extended Data Fig. 2). **(e)** Serial dilutions of WT, *cis1Δ, msp1Δ* and *cis1Δ msp1Δ* were grown on media containing glucose or glycerol as carbon source at 30°C. **(f)** Cell extracts of WT, *cis1Δ, msp1Δ* and *cis1Δ msp1Δ* were analysed by immunoblotting using the indicated antisera. p: precursor; m: mature. **(g)** WT, *cis1Δ, msp1Δ* and *cis1Δ msp1Δ* cells were treated with CCCP to deplete the membrane potential. Where indicated, cells were incubated with cycloheximide (CHX) for the indicated time points. Cell extracts were analysed by immunoblotting with the indicated antisera. p: precursor; m: mature.

Altogether, Msp1 and Ubx2 bind constitutively to the TOM complex to monitor protein import.

### Msp1 promotes degradation of different types of precursor proteins

Msp1 extracts precursor proteins with a bipartite presequence like Cox5a from the TOM complex^29^. A bipartite presequence contains in addition to the presequence a hydrophobic inner membrane sorting sequence. We wondered whether Msp1 also clears translocation-stalled precursor proteins from the TOM complex that lack such a hydrophobic sorting signal and are substrates of the mitoTAD pathway^24^. To address this question, we impaired protein import in *msp1Δ* by the deletion of *PAM17*. Pam17 is the only non-essential subunit of the PAM module^44^. Loss of Pam17 impairs protein transport into the matrix, leading to accumulation of non-imported precursor proteins^44,45^. Previously, we used this approach to uncover a role of mitoTAD components in the removal of precursor proteins such as the Mdj1 precursor that is transported into the matrix^24^. The *msp1Δ pam17Δ* strain displays a mild synthetic growth defect and a mild accumulation of the Mdj1 precursor proteins compared to the single deletion strains (Figs. 4a and b). Supporting this finding, the precursor of the matrix protein Hsp60 accumulates in *msp1*Δ strains when combined with deletion mutants of the TOM subunits^37^ and loss of the mammalian homolog ATAD1 leads to accumulation of the precursor of the ornithin-transcarbamoylase^28^. Conversely, we asked whether Ubx2 is involved in the removal of precursor proteins with a bipartite presequence such as Cox5a. We overexpressed Cox5a_GFP_, which arrests in the TOM-TIM23 supercomplex (Extended Data Fig. 3). In the absence of Msp1 or Ubx2, the precursor of Cox5a_GFP_ accumulates compared to wild-type cells (Fig. 4c), indicating that its removal could be impaired. To confirm this idea, we overexpressed Cox5a_GFP_ and monitored its degradation over time by blocking protein synthesis with cycloheximide. The degradation of the Cox5a_GFP_ precursor is delayed in *msp1Δ* and *ubx2Δ* (Fig. 4d). Moreover, we found that overexpression of Cox5a_GFP_ impairs import of Ilv2, leading to the accumulation of its precursor, which was also not properly removed in the mutant cells (Fig. 4d).

**Figure 4.**
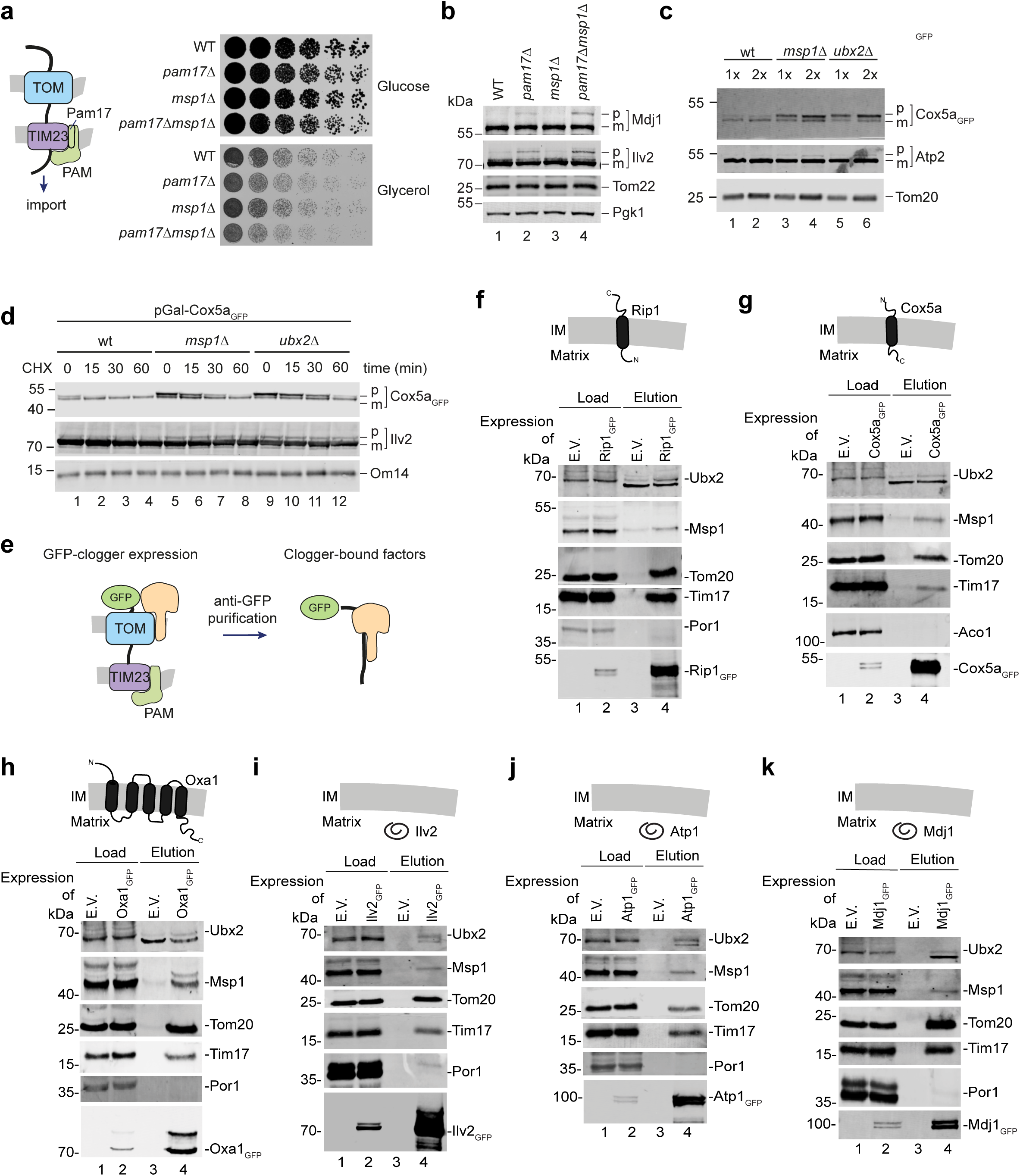
Msp1 and Ubx2 bind different precursor proteins. **(a)** Left: Scheme of the import motor PAM at the TIM23 translocase. Right: Serial dilutions of wild-type (WT), *pam17Δ*, *msp1Δ* and *pam17Δ msp1Δ* were grown on media containing the indicated carbon source at 24°C. **(b)** Cell extracts of WT, *pam17Δ*, *msp1Δ* and *pam17Δ msp1Δ* were analysed by immunoblotting using the indicated antisera. p: precursor; m: mature. **(c)** Cell extracts of WT, *msp1Δ* and *ubx2Δ* expressing Cox5a_GFP_ were analysed by immunoblotting using the indicated antisera. p: precursor; m: mature. **(d)** WT, *msp1Δ* and *ubx2Δ* expressing Cox5a_GFP_ were treated with cycloheximide (CHX) for the indicated time points. Cell extracts were then analysed by immunoblotting with the indicated antisera. p: precursor; m: mature. **(e)** Scheme of the experimental setup to test the interaction of different GFP-tagged precursor proteins with the translocases and quality control factors. **(f) – (k)** WT cells expressing Cox5a_GFP_ (f), Rip1_GFP_ (g), Oxa1_GFP_ (h), Ilv2_GFP_ (i), Atp1_GFP_ (j) or Mdj1_GFP_ (k) and the corresponding empty vector were subjected to affinity purification via the GFP-tag and analysed by immunoblotting using the indicated antisera. Load 0.2 %, elution 100 %.

These observations indicate that Msp1 and Ubx2 have overlapping substrate specificities. To address this possibility, we wondered whether Msp1 and Ubx2 bind to different types of precursor proteins. We overexpressed GFP-fusion constructs of the matrix proteins Mdj1, Ilv2, and Atp2 and of the inner membrane proteins Rip1, Oxa1 and Cox5a (Fig. 4e). Folding of the GFP-domain before the import leads to the arrest of a fraction of the precursor proteins in the TOM-TIM23 supercomplex. We performed affinity purification of these precursor proteins via the GFP-tag and found co-purification of Tom22 of the TOM complex and Tim17 of the TIM23 complex, confirming that the precursor proteins partially arrest in the TOM-TIM23 complex (Figs. 4f-k). Cox5a_GFP_ has a hydrophobic sorting signal and is laterally released from the TIM23 complex, leading to less co-purification of Tim17 (Fig. 4f) compared to other precursor proteins. The other inner membrane proteins Rip1 and Oxa1 are first transported into the matrix before insertion into the inner membrane^46,47^. We also detected Msp1 and Ubx2 in the elution fraction in all pulldowns, indicating that both factors are involved in the removal of various types of precursor proteins (Figs. 4f-k).

We conclude that the central components of mitoCPR and mitoTAD pathways, Msp1 and Ubx2, have overlapping substrate specificities.

### Ubiquitylation of mitochondrial proteins

Ubiquitylation plays an important role in the removal of non-imported precursor proteins^24,39–41^. However, the detection of ubiquitylated proteins is challenging since these proteins are only present for a short time window and rapidly removed. To obtain a comprehensive overview about the ubiquitylated mitochondrial precursor proteins, we co-expressed His-tagged ubiquitin to purify ubiquitylated proteins under denaturing conditions^41^. To enrich ubiquitylated non-imported mitochondrial precursor proteins in the cell, we combined three different strategies (Fig. 5a). First, we deleted *PAM17* to reduce import capacity of mitochondria. Second, we deleted *UBP16* to block deubiquitylation of precursor proteins^41^. Third, we deleted *PDR5* to allow the efficient uptake of the proteasomal inhibitor MG132 to block the degradation of ubiquitylated precursor proteins^41^. We performed affinity purification of His-tagged ubiquitin under these conditions and studied the ubiquitylated proteins by mass spectrometry (Fig. 5b and Supplementary Table 3). Remarkably, we could detect more than 400 ubiquitylated mitochondrial proteins, demonstrating the central role of ubiquitylation in the quality control of mitochondrial proteins. We obtained several lines of evidence that non-imported mitochondrial precursor proteins are ubiquitylated. First, the vast majority of the ubiquitylated mitochondrial proteins are destined for transport into the inner membrane or matrix (Fig. 5c). Second, we could detect a large number of presequence-containing proteins to be ubiquitylated (Fig. 5d). Finally, to demonstrate that particular non-imported mitochondrial proteins are ubiquitylated, we performed a ubiquitin pulldown in cells containing Pam17. When we compared the ubiquitylated proteins in the absence or presence of Pam17, we found that in particularly the ubiquitylation of proteins of the presequence pathway is increased when the import is impaired (Fig. 5e and Supplementary Table 3). We conclude that ubiquitylation plays a central role in the removal of non-imported mitochondrial precursor proteins.

**Figure 5.**
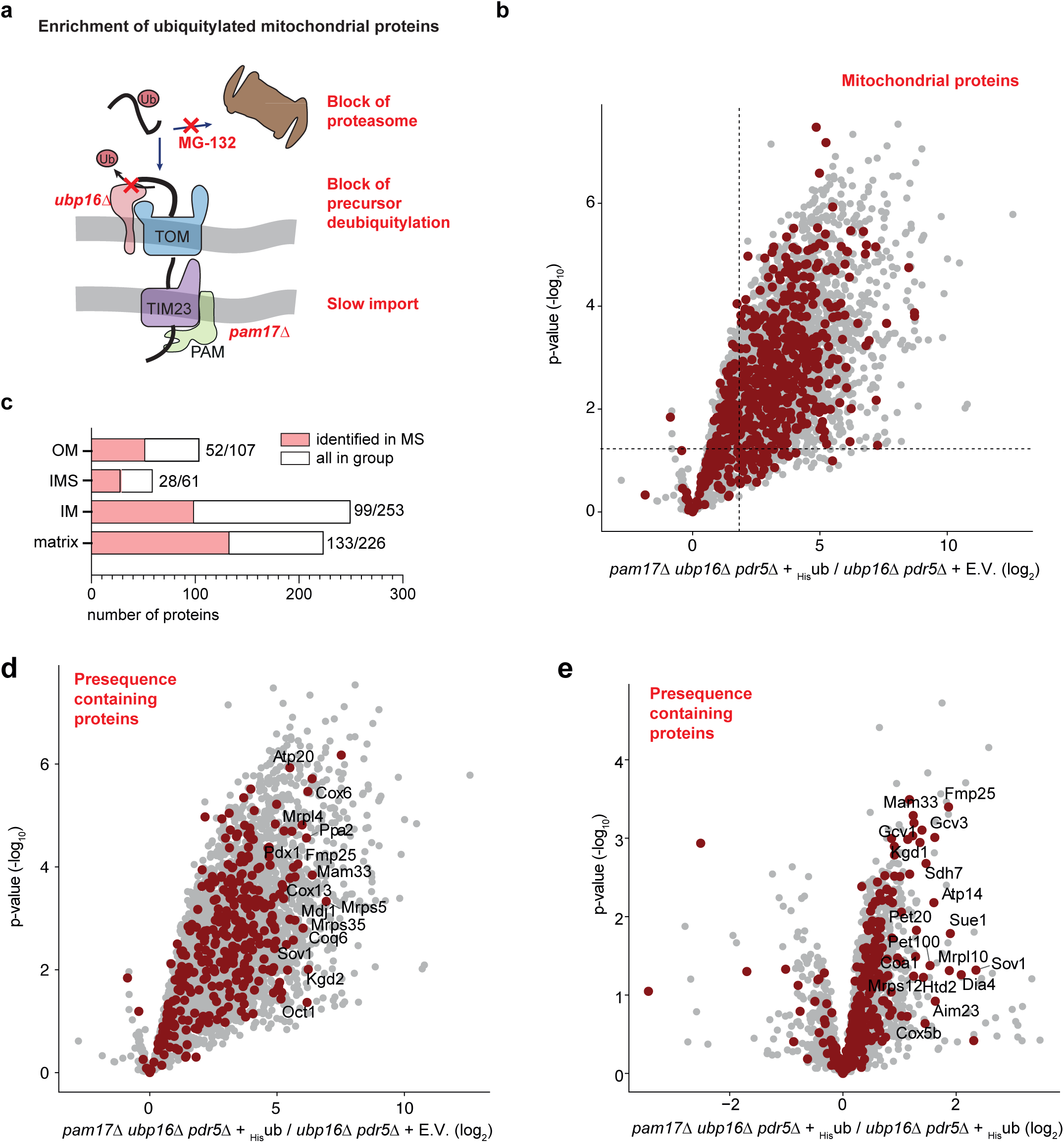
Impaired import triggers ubiquitylation of mitochondrial precursor proteins. **(a)** Scheme displaying the experimental strategy used to enrich ubiquitylated mitochondrial proteins for identification by mass spectrometry. **(b)** Ubiquitylated proteins were enriched by purification via His-tagged ubiquitin from *pam17Δ ubp16Δ pdr5Δ* cells treated with the proteasome inhibitor MG132 under denaturing conditions and compared to *ubp16Δ pdr5Δ* cells not expressing _His_ubiquitin. Four independent replicates were analysed by mass spectrometry. The log_2_ (fold change) of *pam17Δ ubp16Δ pdr5Δ* + _His_ubiquitin versus *pam17Δ ubp16Δ pdr5Δ* expressing the empty vector (E.V.) is plotted against the -log_10_ (p-value). 414 mitochondrial proteins were enriched at least 4-fold over the control with a p-value < 0.05 and were categorized as ubiquitylated mitochondrial proteins. Red dots, mitochondrial proteins. **(c)** Submitochondrial localization of the ubiquitylated proteins identified in (b). The mitochondrial subcompartments were assigned to the proteins according to Morgenstern et al., 2017. **(d)** Data as in (b) with all presequence-containing proteins identified by mass spectrometry marked in red. **(e)** Comparison of ubiquitylated proteins identified by mass spectrometry between *pam17Δ ubp16Δ pdr5*Δ and *ubp16*Δ *pdr5*Δ cells expressing _His_ubiquitin. Presequence-containing proteins are shown in red.

### Msp1 facilitates ubiquitylation of mitochondrial precursor proteins

We found that many non-imported mitochondrial precursor proteins are ubiquitylated. Therefore, we asked if either Ubx2 or Msp1 affect the ubiquitylation of these mitochondrial precursor proteins. We selected the model substrate Cox5a_GFP_, which arrests in the TOM-TIM23 translocon and is bound by both Msp1 and Ubx2 (Fig. 4g). Cox5a_GFP_ was co-expressed with His-tagged ubiquitin in *msp1*Δ and *ubx2*Δ strains followed by affinity purification (Fig. 6a). We detected an increase of ubiquitylated Cox5a_GFP_ in *ubx2*Δ compared to wild-type (Fig. 6a, lanes 7 and 8), indicating that Ubx2 promotes the removal of ubiquitylated Cox5a_GFP_. A similar function of Ubx2 was reported in the ERAD pathway^35,36^. Unexpectedly, we detected a decrease of ubiquitylated Cox5a_GFP_ in the absence of Msp1 (Fig. 6a, lane 9). Similarly, we found that the ubiquitylation of the expressed *b_2_*-DHFR was also decreased in the absence of Msp1 (Fig. 6b). As control, the overall ubiquitylation of proteins was not affected in the mutant strains as shown by immunodetection of all ubiquitylated proteins (Fig 6b). Furthermore, impaired protein import in *pam17*Δ increased the ubiquitylation of overexpressed *b*_2_-DHFR, which is not increased in a *msp1Δ pam17*Δ double deletion strain (Fig. 6b). Thus, the data reveal that Msp1 promotes ubiquitylation of two translocation-arrested precursor proteins.

**Figure 6.**
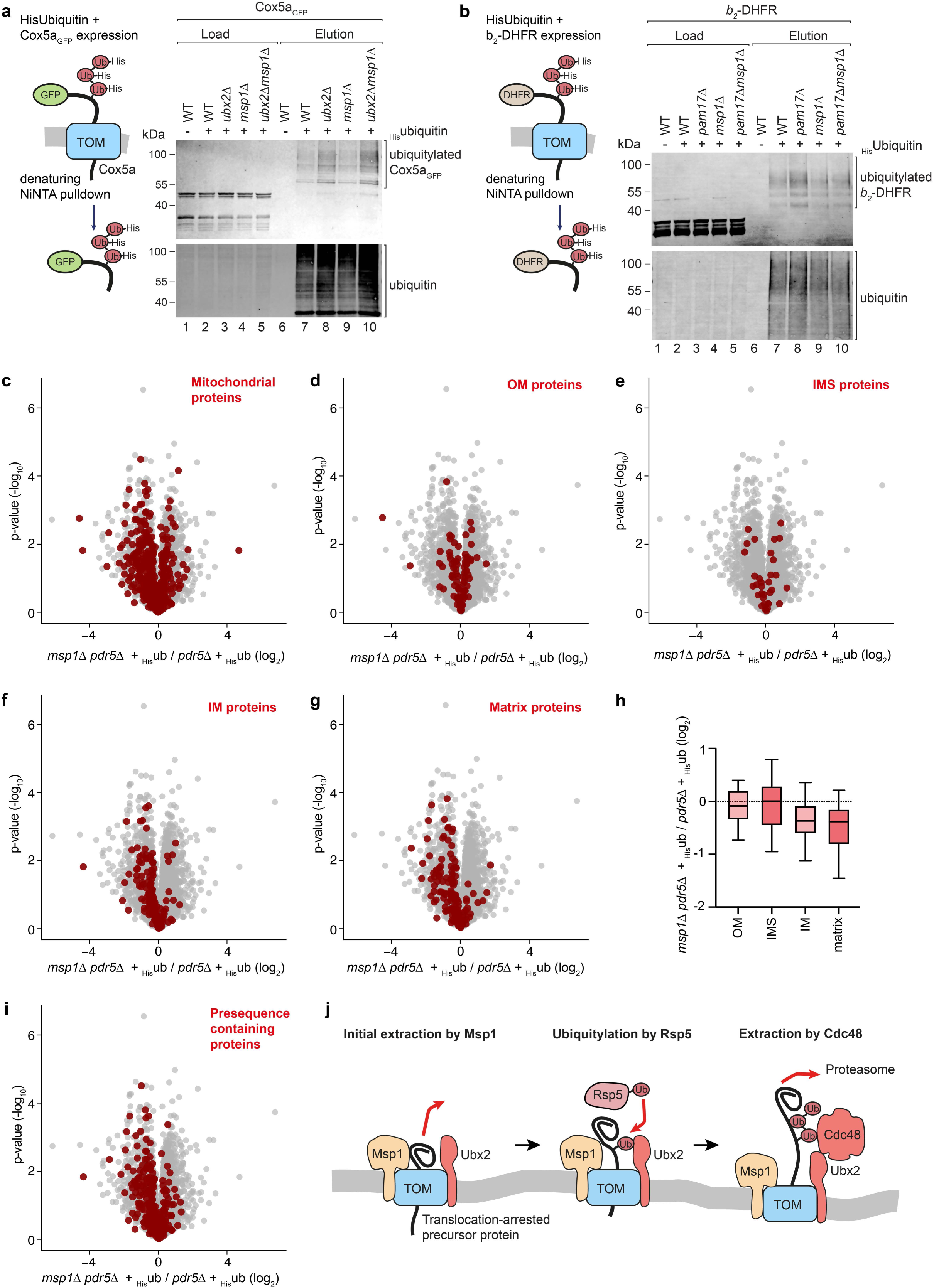
Msp1 facilitates ubiquitylation of presequence-containing proteins. **(a)** Purification of _His_ubiquitin under denaturing conditions from wild-type (WT), *ubx2Δ*, *msp1Δ* and *msp1Δ ubx2Δ* expressing Cox5a_GFP_. Load (0.4 %) and elution (100 %) fractions were analysed by immunoblotting with the indicated antisera. **(b)** Purification of expressed His-tagged ubiquitin under denaturing conditions from WT, *pam17Δ*, *msp1Δ* and *msp1Δ pam17Δ* expressing cytochrome *b_2_*(1-84)-fused to DHFR followed by the heme binding domain of cytochrome *b*_2_ (*b*_2_-DHFR). Load (0.4 %) and elution (100 %) fractions were analysed by immunoblotting with the indicated antisera. **(c) – (g)** Denaturing purifications of _His_ubiquitin from WT and *msp1Δ* cells expressing His-tagged _His_ubiquitin or WT cells containing the corresponding empty vector (E.V.) treated with CCCP and MG132 were analysed by mass spectrometry. Dataset was filtered for proteins at least 4-fold enriched in *msp1Δ* or WT cells expressing _His_ubiquitin compared to the empty vector (E.V.) control. The abundance of the remaining proteins was compared between WT and *msp1Δ* cells expressing _His_ubiquitin. Log_2_ (fold change) of four independent replicates is plotted against the -log_10_ (p-value). Highlighted in red are mitochondrial proteins (c), proteins of the outer membrane (OM) (d), intermembrane space proteins (IMS) (e), inner membrane proteins (IM) (f) and matrix proteins (g). The mitochondrial subcompartments and presequence containing proteins were assigned according to Morgenstern et al., 2017. **(h)** Box plot of median log_2_ (fold change) between WT and *msp1Δ* cells of OM, IM, IMS and matrix proteins detected in the His-ubiquitin pulldown. **(i)** Volcano plot of _His_ubiquitin pulldown as in (c)-(g). Log_2_ (fold change) of four independent replicates is plotted against the -log_10_ (p-value). Highlighted in red are mitochondrial precursor proteins that contain a cleavable presequence. **(j)** Model depicting the cooperation of Msp1 and Ubx2-Cdc48 in the mitoTAD pathway.

We wondered whether Msp1 generally affects ubiquitylation of non-imported mitochondrial precursor proteins. We performed affinity purification via His-tagged ubiquitin in wild-type and *msp1Δ* cells in the presence of CCCP to inhibit mitochondrial import and of MG132 to block degradation of the ubiquitylated proteins. When we analysed the samples by mass spectrometry, we found that the ubiquitylation of mitochondrial proteins was overall decreased (Fig. 6c and Supplementary Table 4). Primarily, ubiquitylation of inner membrane and matrix proteins was reduced in *msp1*Δ, whereas the ubiquitylation of most outer membrane and intermembrane space proteins remained largely unchanged (Figs. 6d-h). Consequently, the ubiquitylation of presequence-containing proteins was decreased (Figs. 6i).

Taken together, our data demonstrate that Ubx2 and Msp1 constitutively monitor protein entry into mitochondria and extract precursor proteins for proteasomal degradation. Msp1 extracts precursor proteins to facilitate their ubiquitylation. In contrast, Ubx2 recruits Cdc48 to complete the extraction of ubiquitylated precursor proteins to deliver them for proteasomal degradation (Figure 6j). The parallel loss Msp1 and Ubx2 leads to impaired removal of non-imported precursor proteins and strongly affects cell viability. Thus, Msp1 and Ubx2-recruited Cdc48 have overlapping functions and substrates to ensure quality control of protein entry at the TOM complex.

## DISCUSSION

Quality control mechanisms at the TOM complex are required to remove translocation-stalled precursor proteins and therefore to ensure full protein import capacity. The mitoTAD and mitoCPR pathways were considered to function independently. According to this view, mitoTAD continuously monitors protein transport at the TOM complex, while the mitoCPR pathway is activated upon import stress^10,11,13–15,24,29^. Our findings change this view. We established that the central subunits of these pathways, Ubx2-Cdc48 and Msp1, cooperate to clear precursor proteins from the TOM complex. Parallel loss of Ubx2 and Msp1 strongly affected cell viability, indicating that both factors have partially overlapping functions. Both proteins associate with the TOM complex independently of stress and bind to various translocation-arrested precursor proteins. Finally, both factors promote the degradation of precursor proteins by the ubiquitin-proteasome system. While Msp1 extracts precursor proteins from the TOM complex to facilitate their ubiquitylation, Ubx2-Cdc48 clears these ubiquitylated precursor proteins to allow their degradation (Figure 6j). We conclude that Ubx2-Cdc48 and Msp1 monitor the TOM complex under both stress and non-stress conditions to remove translocation-arrested precursor proteins. Thus, two AAA-ATPases, Msp1 and Ubx2-recruited Cdc48, ensure the extraction of precursor proteins from the TOM complex.

We found a central role of ubiquitylation in the removal of non-imported mitochondrial proteins. High-throughput studies have revealed that not only many outer membrane proteins but also some inner membrane and matrix proteins can be ubiquitylated^39,48^. Upon impaired protein import and inhibited protein degradation, we could detect several hundred ubiquitylated mitochondrial proteins, including many inner membrane and matrix proteins.

E3-ubiquitin ligases and deubiquitylases control the cycle of ubiquitylation and deubiquitylation of mitochondrial precursor proteins. Whether ubiquitylation is involved in the mitoTAD pathway remained unknown. Based on our observations, we propose that Msp1 mediates the extraction of at least a portion of the precursor protein to expose a site for ubiquitylation. The ubiquitylation site may be shielded when the protein is still in the translocation channel, explaining why Msp1 promotes the ubiquitylation of precursor proteins. Ubx2 recognizes the ubiquitylated protein via its UBA domain^24^ and recruits via its UBX domain the cytosolic Cdc48, which completes the extraction of the precursor protein. Thus, the Ubx2-Cdc48 complex primarily acts on ubiquitylated proteins, while Msp1 particularly extracts non-ubiquitylated precursor proteins. However, Msp1 and Ubx2-Cdc48 are functionally redundant as the single deletion of neither *UBX2* nor *MSP1* compromises protein quality control at the TOM complex. Only the loss of both proteins blocks removal of precursor proteins and is lethal upon import stress.

Msp1 and Ubx2 also remove faulty proteins from the outer membrane^49–56^. Here, Msp1 is particularly important to clear mislocalized and orphaned proteins^49–55^, while Ubx2 promotes the removal of some misfolded proteins^57,58^. Mislocalized proteins are not directly delivered for degradation, but can first be targeted to the endoplasmic reticulum^52^. Whether both factors cooperate in the quality control of these different types of proteins remains to be established. A recent study revealed that Msp1 and Ubx2-Cdc48 are also involved in the removal of translocation-arrested β-barrel precursor proteins from the outer mitochondrial membrane^56^. All these examples reveal the central role of these two AAA-ATPases in maintaining mitochondrial proteostasis.

Altogether, we have established that two AAA-ATPases, the Ubx2-recruited Cdc48 and Msp1, cooperate in the extraction of translocation-stalled mitochondrial precursor proteins from the TOM complex for their degradation by the ubiquitin-proteasome system. This quality control mechanism is critical to maintain protein import into mitochondria and is therefore essential for mitochondrial biogenesis and function.

## MATERIALS AND METHODS

### Yeast and growth conditions

The yeast strains used in this study are listed in the Supplementary Table 5. The open reading frames of *MSP1* and *UBX2* in the BY4741 background were replaced by introduction of a kanMX4, natNT2, hphNT1 or HIS3MX6 selection marker as indicated. The strains were verified by the missing immunosignal in cellular lysate for Msp1 and Ubx2, respectively. *TOM40* was chromosomally fused with a triple HA-tag in different mutant strains as described^24^. The open reading frames of *COX5a*, *ILV2*, *RIP1*, *OXA1*, *MDJ1* and *ATP1* were integrated in a p415 plasmid to fuse them to a GFP. The expression is under control of the GAL1 promoter, allowing a strong expression of all constructs in yeast. The genetic information of His-tagged ubiquitin was integrated into a pRS416 plasmid and transformed into different yeast strains to allow purification of ubiquitylated proteins (Supplementary Table 6)^41^. Yeast strains were grown on full medium containing 2% (w/v) glucose or 3% (w/v) glycerol at 24-30°C. In case yeast cells were transformed with a plasmid, cells were grown on selective medium (0,67% (w/v) yeast nitrogen base, 0,07% (w/v) amino acid mixture lacking uracil or leucine or both) containing 2% (w/v) glucose as carbon source. For expression of plasmid-encoded precursor proteins under the control of the GAL promoter, cells were grown in selective medium containing 2 % (w/v) raffinose as carbon source till they reached the exponential growth phase. Expression by the *GAL1* promoter was induced by addition of 2% (w/v) galactose.

### Preparation of cell extracts

Cell extracts were prepared by post-alkaline extraction following a published procedure^59^. Cells were grown to the exponential growth phase and harvested by centrifugation (2,500*g*, 5 min, 20°C) and washed with water. The pellet was resuspended in 0.2 M NaOH and incubated for 5 min at 20°C. Subsequently, cells were pelleted (2,500*g*, 5 min, 20°C) and lysed with HU sample buffer for 10 min at 65°C.

### Isolation of mitochondria

Mitochondria were isolated by differential centrifugation^60^. Yeast cultures were grown until an early logarithmic growth phase. Cells were harvested (3,000*g*, 8 min, 20°C), washed with water and resuspended in dithiothreitol (DTT) buffer (0.1 M Tris/H_2_SO_4_ pH 9.4 and 10 mM DTT). Cells were incubated in DTT buffer for 30 min at 30°C under constant shaking. Yeast cells were re-isolated, washed and resuspended in zymolyase buffer (1.2 M sorbitol, 20 mM KPi pH 7.4) containing zymolyase (5 mg/g cells) and incubated for 45 min at 30°C under constant shaking to open the cell wall. The obtained spheroplasts were re-isolated and homogenized in ice cold homogenisation buffer (0.6 M sorbitol, 10 mM Tris/HCl pH 7.4, 1 mM ethylenediaminetetraacetic acid (EDTA), 1 mM phenylmethylsulfonylfluoride (PMSF), 0.2% (w/v) bovine serum albumin). Subsequently, unbroken cells and cell debris were removed (3,000*g*, 5 min, 4°C) and a mitochondrial pellet was isolated by centrifugation (17,000*g*, 10 min, 4°C). Mitochondria were washed before they were resuspended in SEM buffer (250 mM sucrose, 10 mM MOPS, 1 mM EDTA). Aliquots were shock frozen in liquid nitrogen and stored at -80°C until further use.

### Analysis of mitochondrial protein complexes

Mitochondrial protein complexes were studied by blue native electrophoresis^61^. The blue native gel was prepared as described^60^. For the sample preparation, mitochondrial pellets were lysed with 1% (w/v) digitonin in lysis buffer (0.1 mM EDTA, 50 mM NaCl, 10% (v/v) glycerol, 20 mM Tris/HCl pH 7.4) for 15 min on ice. Subsequently, the samples were subjected to centrifugation (17,000*g*, 10 min, 4°C) to remove insoluble material. Blue native loading dye was added to the supernatants and protein complexes were separated on a blue native gel.

### Affinity purification of the TOM complex

To purify the TOM complex from yeast or isolated mitochondria, Tom40 was chromosomally fused to a triple HA-tag. For purification out of isolated mitochondria, mitochondria were lysed with 1% (w/v) digitonin in lysis buffer (20 mM Tris/HCl pH 7.4, 50 mM NaCl, 10% (w/v) glycerol, 0.1mM EDTA containing 1 mM PMSF, 20 mM N-ethylmaleimide (NEM), HALT protease inhibitor cocktail for 15 min at 4°C under constant rotation. Insoluble material was removed by centrifugation (17,000*g*, 10 min, 4°C) and the mitochondrial lysate was incubated with preequilibrated anti-HA beads (Roche) for 1.5 h at 4°C. The beads were excessively washed with washing buffer (20 mM Tris/HCl pH 7.4, 50 mM NaCl, 10 % (w/v) glycerol) containing 20 mM NEM, HALT protease inhibitor cocktail and 0.1 % (w/v) digitonin. Bound proteins were eluted with HU-buffer (5 % SDS (w/v), 8 M Urea, 1,5 % (w/v) DTT, 1 mM EDTA, 200 mM Tris-HCl pH 6.8, 0.025 % (w/v) bromphenol blue). For pulldowns out of total cell extract, cells corresponding to 200 optical densities at 600 nm (OD_600_) were ruptured by mechanic force after adding silica beads at 4°C using a RETSCH MM400 mill. After removal of beads and unbroken cells by centrifugation (3,000*g*, 5 min, 4°C), purification was continued as described with isolated mitochondria.

### Affinity purification of precursor proteins

GFP-fused mitochondrial proteins were overproduced in different yeast strains. Proteins were co-purified utilizing GFP-trap beads (ChromoTek). Cell extracts were prepared as described above. Cell extracts were lysed with 1% (w/v) digitonin lysis buffer for 15 min at 4°C under constant rotation. Insoluble materials were removed by centrifugation (17,000*g*, 10 min, 4°C) and samples were incubated with anti-GFP agarose for 1.5 h at 4°C under constant rotation. After excessive washing with wash buffer, the proteins were eluted by incubation with HU-buffer at 65°C for 10 min.

### Protein stability assay

Exponentially growing yeast cell cultures were treated with 100 µg/ml cycloheximide to block cytosolic protein synthesis^41^. 2.5 OD_600_ were collected at different timepoints and whole cell extracts were prepared by incubation with 200 mM NaOH for 5 min at 20°C followed by centrifugation (17,000*g*, 10 min, 4°C). Pellets were resuspended in HU buffer and incubated at 65°C for 10 min. For the depletion of the membrane potential, cells were treated for 1 h with 20 µM CCCP prior the stability assay.

### Quantitative PCR

For quantification of the relative transcript levels of exponentially grown yeast cells expressing a *b*_2_-DHFR or the empty vector control, RNA was isolated using the NucleoSpin RNA isolation kit (Macherey-Nagel), followed by DNase I treatment and cDNA synthesis (320 ng/20 µl) using RevertAid Reverse Transcriptase (Thermo Fisher Scientific) and oligo-d(T)_23_ VN primers (New England Biolabs). In order to ensure the removal of any DNA contamination prior to qPCR, RT-PCR on cDNA and DNase-treated RNA using the respective qPCR primer pair was executed. RT-qPCR was performed using a 7300 Real-Time PCR System together with the Evagreen master mix (Biobudget) and an amount of cDNA corresponding to 10 ng RNA. As reference gene, the transcription factor A (TFA2) gene was used. The study was performed using three biological replicates with three technical replicates each. The primers used are listed in Supplementary Table 7. The annealing temperature was set to 60 °C and the elongation time was set to 20 seconds in 40 amplification cycles. The validation of clean runs was facilitated by the use of the Applied Biosystems 7300/7500/7500 Fast System software. The relative expression levels of the respective genes were calculated using the 2^-ΔΔCt^ method^62^.

### Affinity purification of His-tagged ubiquitin

A plasmid encoding for His-tagged ubiquitin was transformed in different mutant strains to allow expression of His-tagged ubiquitin in yeast cells^63^. To increase the amount of ubiquitylated proteins, *pdr5*Δ cells were treated with the proteasomal inhibitor MG132 (100 µM) for 1h prior affinity purification. The affinity purification of His-tagged ubiquitin was performed as described^41^. For this, 200 OD_600_ of cells were harvested and resuspended in buffer A (6 M guanidine HCl, 100 mM NaH_2_PO_4_ and 10 mM Tris pH 7.4) and mechanically ruptured using silica beads for 5 min at 4°C. After removal of unbroken cells and beads by centrifugation (3,000*g*, 5 min, 4 °C), 0.05% (v/v) Tween 20 and 20 mM imidazole was added. After removal of insoluble material by centrifugation (17,000*g*, 10 min, 4°C), the supernatant was incubated with Ni^2+^-NTA agarose beads (Qiagen) for 1 h 30 min at 4°C under constant rotation. Beads were then washed with buffer A containing 0.05% (v/v) Tween 20 and 20 mM imidazole, followed washing steps with buffer C (8 M urea, 100 mM NaH_2_PO_4_ and 10 mM Tris pH 6.8). Bound proteins were eluted with HU buffer at 65°C for 10 min.

### Sample preparation for LC-MS/MS

The protein samples of the affinity-purifications of Tom40_HA_ and His-tagged ubiquitin from yeast were prepared for LC-MS/MS by in-gel digestion (IGD) or were processed using single-pot, solid-phase-enhanced sample preparation (SP3)^64^. For IGD, proteins were briefly separated by SDS-PAGE and gel lanes were cut into 3-4 slices. Gel slices were destained in 50 mM ammonium bicarbonate/acetonitrile (1:1, v/v), reduced with 20 mM DTT for 20 min at 60°C, alkylated with 40 mM acrylamide for 30 min at room temperature in the dark, and digested overnight at 37°C with sequencing-grade trypsin (Promega GmbH, Walldorf, Germany) at 5 ng/µL in 50 mM ammonium bicarbonate. Peptides were extracted sequentially with 50% (v/v) and 100% (v/v) acetonitrile, dried, and reconstituted in 0.1% (v/v) formic acid. For other samples were prepared with an in-solution protocol (single-pot, solid-phase-enhanced sample preparation (SP3)^64^: proteins were reduced with 20 mM DTT, alkylated with 40 mM iodoacetamide, subjected to SP3 cleanup with a mixture of hydrophilic carboxylate-coated magnetic beads (Sera-Mag SpeedBeads), and digested with trypsin at an enzyme-to-protein ratio of 1:25 (w/w) in 50 mM Triethylammonium bicarbonate. 10 µg of peptides were further desalted with C18 ZipTips (Merck Millipore, Darmstadt, Germany) to ensure complete removal of beads. Approximately 0.5-1 µg of peptide was injected per LC-MS/MS analysis.

### NanoLC-MS/MS and data-independent acquisition

Samples were loaded onto a self-packed analytical C18 column (400 mm × 75 µm inner diameter; 3 µm ReproSil-Pur 120 C18-AQ, Dr. Maisch). Peptides were separated using a gradient from 5% to 35% solvent B (90% acetonitrile, 0.1% formic acid) and analyzed after nanospray ionization with an Orbitrap Fusion Lumos mass spectrometer (ThermoFisher Scientific). Peptides from the initial Tom40-HA affinity purification were separated during a 90 min gradient and data were acquired in data-dependent mode (DDA) after higher energy collision (HCD) induced dissociation with detection in the Orbitrap.

All other samples were analyzed in data-independent acquisition (DIA) mode with a gradient length of 120 min. Tom40-HA-cis1Δ and HisUb samples were analyzed with variable isolation window width HCD fragmentation (stepped normalized collision energies of 23, 28, and 33). Normalized MS2 AGC target was 1000% and the maximum injection time was 54 ms, spectra were acquired in profile mode. Narrower windows were used in regions of higher precursor density and broader windows at higher m/z values. pam17 samples were acquired with isolation in 25 fixed windows of 24 m/z width. For enhanced library creation aliquots of each sample were pooled and measured with an extended gradient DDA method.

### MS data processing and protein quantification

All data were searched against the *Saccharomyces cerevisiae* UniProt proteome supplemented with the contaminants database by Frankenfield et al. [Frankenfield, A.M., et al., Protein Contaminants Matter: Building Universal Protein Contaminant Libraries for DDA and DIA Proteomics. J Proteome Res, 2022. 21(9): p. 2104-2113.].

DDA data were processed in Proteome Discoverer 2.5.0.400 (Thermo Fisher Scientific) using Mascot Server 2.8.2 for protein identification. DIA data were processed with DIA-NN 1.8.1., Spectronaut 18.5 or Chimerys 4.0.25 (from Proteome Discoverer 3.2.0.450). MS2 tolerance was always set to 20 ppm, oxidation of methionine was considered as dynamic modification, alkylation of cysteine as fixed modification. False discovery rates were controlled at 1% at the peptide and protein levels, and proteins supported by at least two unique peptides were retained for downstream analysis.

### Protein-level statistical analysis

Protein abundances exported from the data processing software were analysed in Perseus (version 4.5.1). Intensities were log2-transformed. Proteins were retained when at least two valid values were present in at least one comparison group when three replicates were measured (Tom40_HA_ purified from mitochondria) or when at least three valid values were present in at least one comparison group (all other measurements). Missing values were imputed from a random distribution with a downshift of 1.8 and 0.3 width. Differential abundance was calculated as the difference between group means and assessed using two-sided Welch’s t-tests. For identification of ubiquitylated proteins, values of His-ubiquitin pulldowns were first compared to the wild-type control. Proteins were defined as ubiquitylated using p < 0.05 and log2FC ≥ 2 as cut-off. For pairwise comparisons of ubiquitylated proteins between *pam17Δ ubp16Δ pdr5Δ* with *ubp16Δ pdr5Δ* or wild-type with *msp1Δ*, the remaining proteins were normalized to the mean abundance of all values.

Data and code availability: The mass spectrometry proteomics data have been deposited to the ProteomeXchange Consortium via the PRIDE partner repository under accession number [PLACEHOLDER].

### Immunoblotting

Proteins and protein complexes were separated by SDS-PAGE or blue native electrophoresis, respectively. Subsequently, proteins were transferred onto a polyvinylidene fluoride (PVDF) membrane using semi-dry Western blotting. Therefore, gels, the PVDF membrane and filter papers were soaked in blotting buffer (20% (v/v) ethanol, 20 mM Tris, 150 mM glycine, 0,02% (w/v) SDS) before Western blotting. The membrane was excessively washed with TBS buffer (20 mM Tris HCl pH 7.4, 150 mM NaCl) and free binding sites were blocked with 5% (w/v) skimmed milk powder or RotiBlock in TBS buffer for overnight at 4°C. We used different primary antisera raised in rabbits for the detection of the proteins (Supplementary Table 8). The specificities of the used antisera were controlled by the absence of immunosignals in cellular or mitochondrial lysates from the respective deletion strains. In case of essential genes, the size shift of the detected band in lysates from cell expressing tagged proteins confirmed the specificity of the immunosignal of the primary antiserum. Secondary antisera with fluorescence labels (IRDye800CW) were used to visualize the signals. The immune signals were detected by using the Odyssey CLx infrared imaging system (Li-Cor). We indicated in the figures by separating white lanes when irrelevant bands have been digitally removed.

## Supporting information

Engelke et al Table S1

Engelke et al Table S2

Engelke et al Table S4

Engelke et al Table S3

## Data and code availability

The mass spectrometry proteomics data have been deposited to the ProteomeXchange Consortium via the jPOST repository. The datasets can be found under the identifier PXD083689 in ProteomeXchange and JPST004895 in JPOST. During the review process, the deposited data can be accessed at https://repository.jpostdb.org/preview/4450353556a9aac6f05a8a using the access key 3301.

## ACKNOWLEDGEMENTS

Work in this study has also been performed in partial fulfilments of the requirements for the doctoral thesis of J.E., N.O., L.P. and K.R.. We thank Ralph Mahlberg, Hannah Scheuch, Jacqueline Schiessl, Lina Letizia and Beatrix Brummer for expert technical assistance. This study was supported by the Deutsche Forschungsgemeinschaft (DFG) (SFB 1218 B11 project ID 269925409, BE 4679/9-1 project ID 528247081, priority program SPP 2453 BE 4679/11-1 project ID 541555098, to T.B.; BE 4679/13-1 project number: 568735694; BR 6283/5-1 project ID 529716110, BR 6283/6-1 project ID 541596792, to F.d.B.) The mass spectrometer of the Core Facility “Analytical Proteomics”, University of Bonn, was funded by the DFG (project ID 386936527).

## AUTHOR CONTRIBUTIONS

Author contributions: J.E., N.O., L.P., K.R., R.H., E.S. and M.S. performed the experiments and analysed the data together with F.d.B. and T.B.. T. B. and F.d.B. designed and supervised the project; T. B., F.d.B. and J.E. designed and prepared the figures. T.B. and F.d.B. wrote the manuscript; all authors discussed results from the experiments and commented on the manuscript.

## Extended Data – Figure legends

**Extended Data Figure 1.**
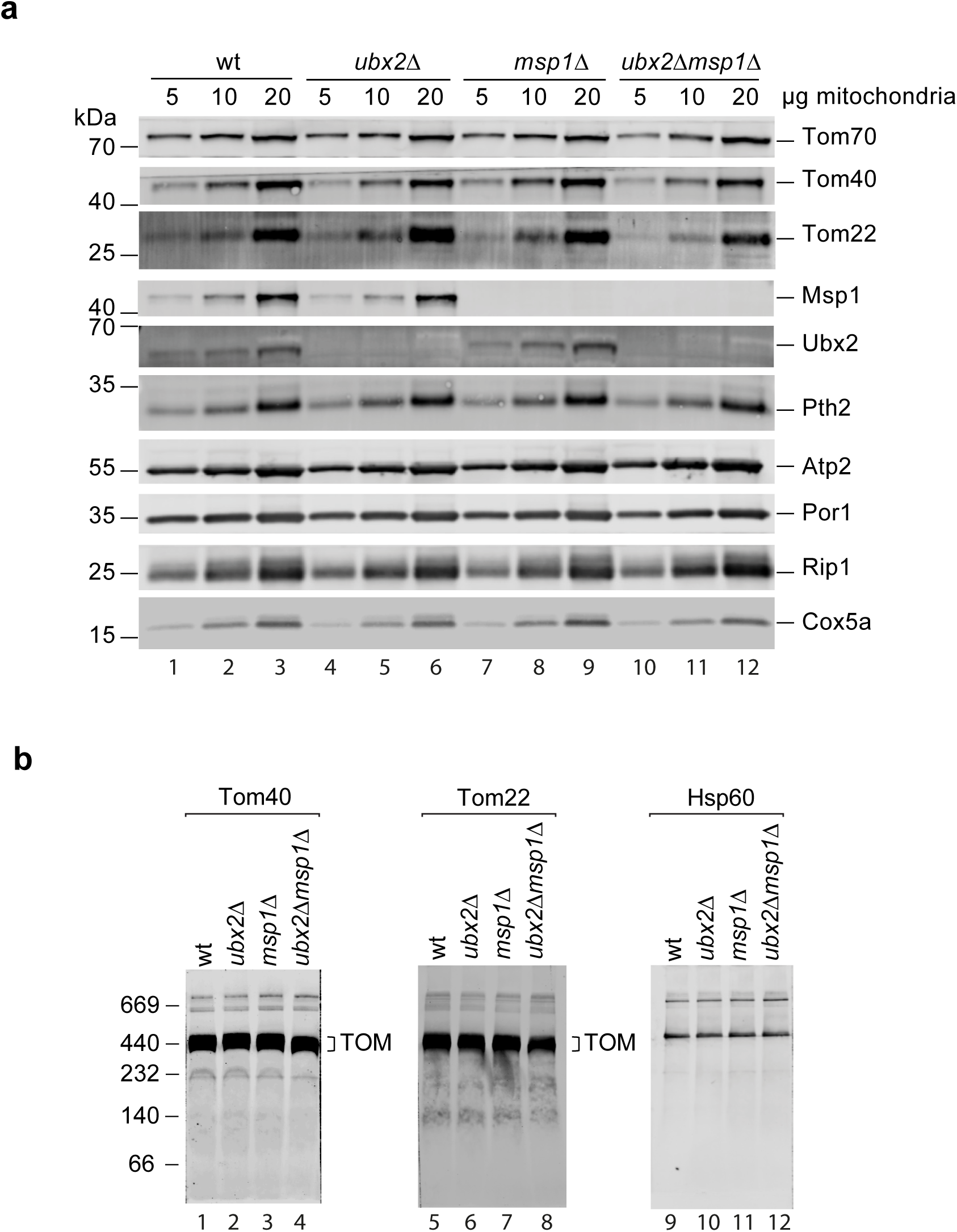
Characterization of *ubx2*Δ, *msp1*Δ and *ubx2Δ msp1Δ* mitochondria. **(a)** Isolated mitochondria from wild-type (WT), *ubx2Δ*, *msp1Δ* and *ubx2Δ msp1Δ* cells were analysed by immunoblotting with the indicated antisera. **(b)** Isolated mitochondria from wild-type (WT), *ubx2Δ*, *msp1Δ* and *ubx2Δ msp1Δ* cells were lysed and protein complexes were analysed by blue native gel electrophoresis followed by immunoblotting with the indicated antisera.

**Extended Data Figure 2.**
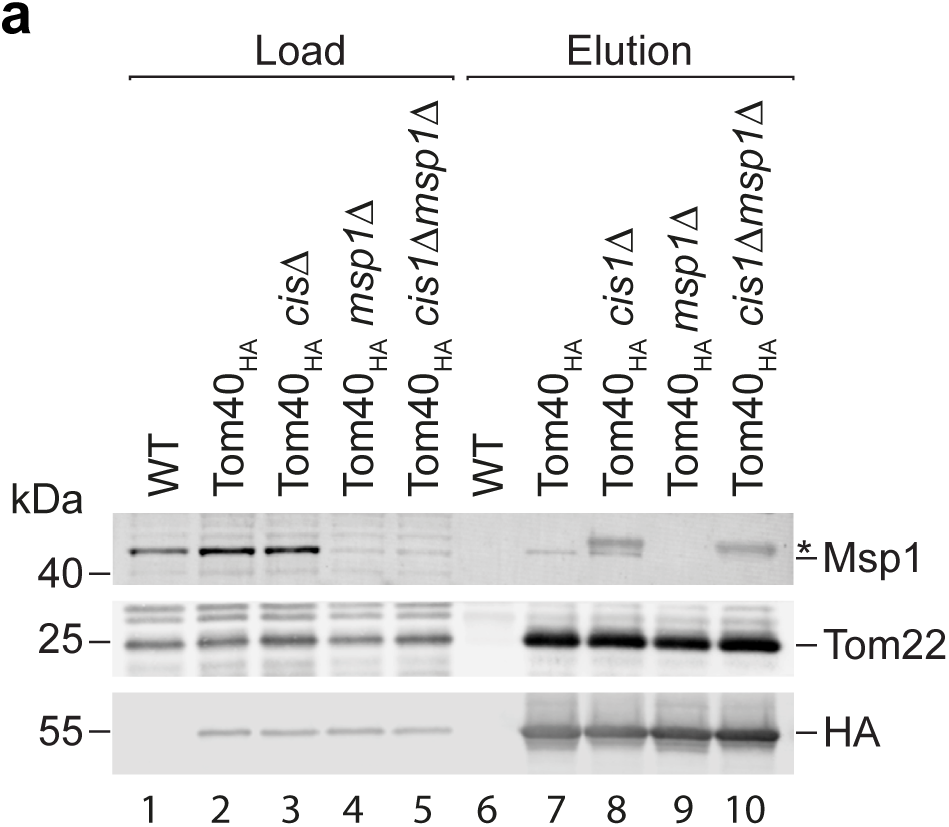
Tom40_HA_ affinity purification in *cis1*Δ. **(a)** Affinity purifications from cell extracts of wild-type (WT), Tom40_HA_ and Tom40_HA_ *cis1*Δ, Tom40_HA_ *msp1*Δ and Tom40_HA_ *cis1Δ msp1*Δ cells were analysed by immunoblotting with the indicated antisera. Load (0.2 %), elution (100 %). Asterisk denotes an unspecific cross-reactive band of the Msp1 antiserum that appears upon deletion of *CIS1*, which is also detected in absence of Msp1 (lane 10).

**Extended Data Figure 3.**
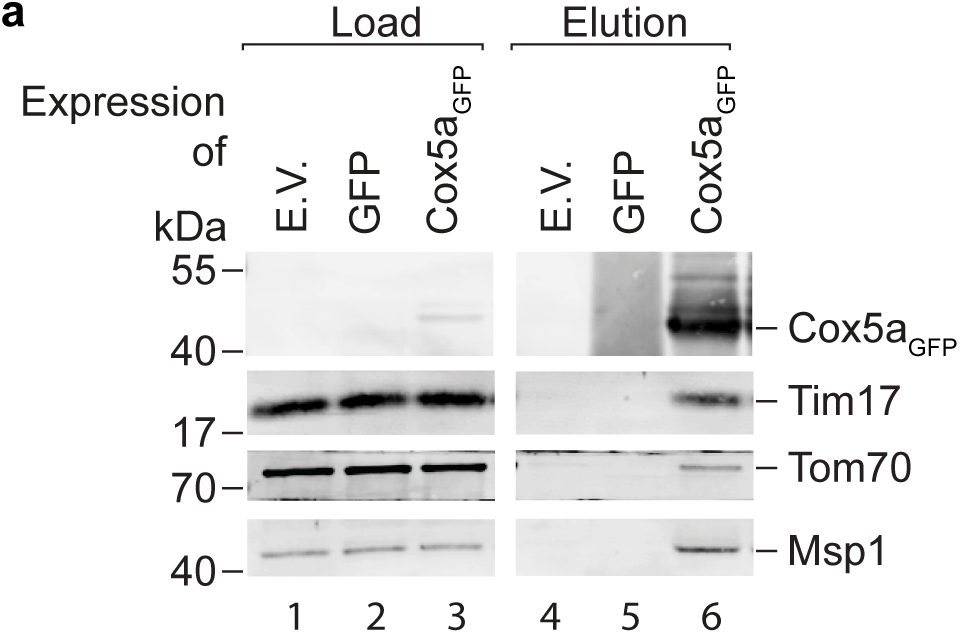
Cox5a_GFP_ partially arrests at the TOM and TIM23 translocases. **(a)** Wild-type (wt) cells expressing Cox5a_GFP_, GFP or the corresponding empty vector were subjected to affinity purification utilizing the GFP-tag. Load (0.2 %) and elution fraction (100 %) were analysed by immunoblotting using the indicated antisera.

## SUPPLEMENTARY INFORMATION

**Supplemental Tables, provided as Excel files:**

**Table S1** | List of proteins identified in the affinity purification of Tom40_HA_ from isolated mitochondria by quantitative MS.

**Table S2** | List of proteins identified in the affinity purification of Tom40_HA_ and Tom40_HA_ *cis1Δ* from cell extracts by quantitative MS.

**Table S3** | List of proteins identified in the affinity purification of His-tagged ubiquitin from *pdr5Δ upb16Δ pam17Δ* and *pdr5Δ upb16Δ* cell extracts by quantitative MS.

**Table S4** | List of proteins identified in the affinity purification of His-tagged ubiquitin from *msp1Δ* cell extracts by quantitative MS.

**Table S5** | Yeast strains used in this study.

**Table S6** | Plasmids used in this study.

**Table S7** | Oligonucleotides used in this study.

**Table S8** | Antisera used in this study.

