## Supplementary Information for "Msp1-dependent extraction promotes ubiquitylation of translocation-stalled mitochondrial precursor proteins"

**Supplemental Tables, provided as Excel files:**

**Table S1** | List of proteins identified in the affinity purification of Tom40$_{HA}$ from isolated mitochondria by quantitative MS.

**Table S2** | List of proteins identified in the affinity purification of Tom40$_{HA}$ and Tom40$_{HA}$ *cis1*Δ from cell extracts by quantitative MS.

**Table S3** | List of proteins identified in the affinity purification of His-tagged ubiquitin from *pdr5*Δ *upb16*Δ *pam17*Δ and *pdr5*Δ *upb16*Δ cell extracts by quantitative MS.

**Table S4** | List of proteins identified in the affinity purification of His-tagged ubiquitin from *msp1*Δ cell extracts by quantitative MS.

**Table S5** | Yeast strains used in this study.

| **Genotype** | **Source** | **Identifier** |
|---|---|---|
| BY4741 (WT) MATa, *his3Δ1*, *leu2Δ0*, *met15Δ0*, *ura3Δ0* | EUROSCARF | TB2 |
| BY4741 *ubx2::kanMX4* | EUROSCARF | TB22 |
| BY4741 *msp1::HIS3MX6* | this study | TB547 |
| BY4741 *msp1::kanMX4, ubx2::hphNT1* | *Ref. 24* | TB604 |
| BY4741 *vms1::kanMX4* | EUROSCARF | TB23 |
| BY4741 *msp1::HIS3MX6, vms1::kanMX4* | this study | TB599 |
| BY4741 *pth2::kanMX4* | EUROSCARF | TB8 |
| BY4741 *msp1::HIS3MX6, pth2::kanMX4* | this study | TB600 |
| BY4741 WT + yep352 | this study | TB713 |
| BY4741 WT + p$b_2$-DHFR | this study | TB714 |
| BY4741 *ubx2::kanMX4* + pDP $b_2$-DHFR | this study | TB715 |
| BY4741 *msp1::HIS3MX6* + pDP $b_2$-DHFR | this study | TB716 |
| BY4741 *msp1::kanMX4, ubx2::hphNT1* +pDP $b_2$-DHFR | this study | TB717 |
| BY4741 WT + pDP-$b_2$Δ-DHFR | this study | TB718 |
| BY4741 *ubx2::kanMX4* + pDP $b_2$Δ-DHFR | this study | TB719 |
| BY4741 *msp1::HIS3MX6* + pDP $b_2$Δ-DHFR | this study | TB720 |
| BY4741 *msp1::kanMX4, ubx2::hphNT1* + pDP $b_2$Δ-DHFR | this study | TB721 |
| BY4741 *tom40::TOM40-3HA-HIS3MX6* | *Ref. 24* | TB4 |
| BY4741 *tom40::TOM40-3HA-HIS3MX6* + pYEp352 | this study | TB722 |
| BY4741 *tom40::TOM40-3HA-HIS3MX6* + pDP $b_2$-DHFR | this study | TB723 |
| BY4741 *tom40::TOM40-3HA-HIS3MX6* + pDP $b_2$Δ-DHFR | this study | TB724 |

| | | |
|---|---|---|
| BY4741 *pth2::kanMX4*, *tom40::TOM40-3HA-HIS3MX6* | *Ref. 41* | TB5 |
| BY4741 *ubx2::kanMX4*, *tom40::TOM40-3HA-HIS3MX6* | *Ref. 24* | TB6 |
| BY4741 *msp1::kanMX4*, *tom40::TOM40-3HA-HIS3MX6* | this study | TB589 |
| BY4741 *cis1::kanMX4*, *tom40::TOM40-3HA-HIS3MX6* | this study | TB588 |
| BY4741 *msp1::hphNT2*, *cis1::kanMX4*, *tom40::TOM40-3HA-HIS3MX6* | this study | TB415 |
| BY4741 *cis1::kanMX4* | EUROSCARF | TB120 |
| BY4741 *cis1::kanMX4*, *ubx2::natNT2* | this study | TB601 |
| BY4741 *pam17::hphNT1* | *Ref. 24* | TB9 |
| BY4741 *pam17::hphNT1*, *msp1::HIS3MX6* | this study | TB592 |
| BY4741 WT + p415 pGAL Cox5a-GFP | this study | TB725 |
| BY4741 *ubx2::kanMX4* + p415 pGAL Cox5a-GFP | this study | TB726 |
| BY4741 *msp1::HIS3MX6* + p415 pGAL Cox5a-GFP | this study | TB727 |
| BY4741 WT + p415 pGAL GFP | this study | TB728 |
| BY4741 WT + p415 pGAL Rip1-GFP | this study | TB729 |
| BY4741 WT + p415 pGAL Oxa1-GFP | this study | TB730 |
| BY4741 WT + p415 pGAL Ilv2-GFP | this study | TB731 |
| BY4741 WT + p415 pGAL Atp1-GFP | this study | TB732 |
| BY4741 WT + p415 pGAL Mdj1-GFP | this study | TB733 |
| BY4741 WT + p415 pGAL | this study | TB734 |
| YPH499 *ubp16::natNT2*, *pdr5::HIS3MX6* + prs415 | this study | TB735 |
| YPH499 *ubp16::natNT2*, *pdr5::HIS3MX6* + HisUbiquitin (LEU2) | this study | TB736 |
| YPH499 *ubp16::natNT2*, *pdr5::HIS3MX6, pam17::hphNT1*, + HisUbiquitin (LEU2) | this study | TB737 |
| BY4741 WT + p415 pGAL Cox5a-GFP + prs416 | this study | TB738 |
| BY4741 WT + p415 pGAL Cox5a-GFP + prs416 HisUbiquitin (URA3) | this study | TB739 |
| BY4741 *ubx2::KanMX4* + p415 pGAL Cox5a-GFP + pRS416 HisUbiquitin (URA3) | this study | TB740 |
| BY4741 *msp1::HIS3MX6* + p415 pGAL Cox5a-GFP + pRS416 HisUbiquitin (URA3) | this study | TB741 |
| BY4741 *msp1::kanMX4, ubx2::hphNT1* + p415 pGAL Cox5a-GFP + pRS416 HisUbiquitin (URA3) | this study | TB742 |
| BY4741 WT + $b_2$-DHFR + prs315 | this study | TB743 |
| BY4741 WT + $b_2$-DHFR + HisUbiquitin (LEU2) | this study | TB744 |
| BY4741 *pam17::hphNT1* + pDP $b_2$-DHFR + pRS415 HisUbiquitin (LEU2) | this study | TB745 |
| BY4741 *msp1::HIS3MX6* + pDP $b_2$-DHFR + pRS415 HisUbiquitin (LEU2) | this study | TB746 |
| BY4741 *msp1::HIS3MX6, pam17::hphNT1* + pDP $b_2$-DHFR + pRS415 HisUbiquitin (LEU2) | this study | TB747 |
| BY4741 *pdr5::natNT2* + pRS415 (LEU2) | this study | TB564 |
| BY4741 *pdr5::natNT2* + pRS415 HisUbiquitin (LEU2) | this study | TB561 |

| | | |
|---|---|---|
| BY4741 *pdr5::natNT2, msp1::HIS3MX6* + pRS415 HisUbiquitin (LEU2) | this study | TB563 |

**Table S6** | Plasmids used in this study.

| **Plasmid** | **Source** | **Identifier** |
|---|---|---|
| pYEp352 | *Ref. 24* | pTB52 |
| pDP-CYB2(1-84)-DHFR-HB(81-181) (referred to as $b_2$-DHFR) | *Ref. 65* | pTB7 |
| pDP-CYB2(1-84)Δ(47-65)-DHFR-HB(81-181) (referred to as $b_2$Δ-DHFR) | *Ref. 65* | pTB8 |
| p415 pGAL | *Ref. 66* | pTB41 |
| p415 pGAL GFP | *Ref. 66* | pTB153 |
| p415 pGAL Cox5a-GFP | this study | pTB164 |
| p415 pGAL Rip1-GFP | this study | pTB158 |
| p415 pGAL Oxa1-GFP | this study | pTB157 |
| p415 pGAL Ilv2-GFP | this study | pTB165 |
| p415 pGAL Atp1-GFP | this study | pTB155 |
| p415 pGAL Mdj1-GFP | this study | pTB156 |
| pRS416 8xHisUbiquitin (URA) | this study | pTB240 |
| pRS415 8xHisUbiquitin (LEU2) | *Ref. 67* | pTB3 |
| pRS315 | *Ref. 67* | pTB4 |
| pRS415 | *Ref. 68* | pTB103 |
| pRS416 | *Ref. 68* | pTB104 |

**Table S7** | Oligonucleotides used in this study.

| **Name** | **Sequence** |
|---|---|
| qPCR CIS1 for | ATCAGTAATTGTCCCATCGGGTTAGTTTC |
| qPCR CIS1 rev | CCTGGGCAGCCTTGAGTAAATCATATC |
| qPCR TFA2 for | CGATTCTTCAAAGTTGCTTTGGGCGAC |
| qPCR TFA2 rev | GTTCCGAAGGGGAATGGACATCGTA |
| Msp1_S1 | AAGGAAGAAGCAAGAACGAAAAGAGATAAGGATTCAAAAGAAAGGAA GCCCAATGCGTACGCTGCAGGTCGAC |
| Msp1_S2 | GATATGATGCGTGAATAAAAAGCTTTCTTCTTTTTTTTCTAATTTTCCTT CCTTAATCGATGAATTCGAGCTCG |
| Tom40 S2 | AAAACAATGATTATTTATTCAACCATAAAAAAGCCAAGGGAAGATTTTC AATCGATGAATTCGAGCTCG |
| Tom40 S3 | AACAAGGTTTAGACGCAGATGGTAACCCATTGCAAGCTCTTCCTCAAT TGCGTACGCTGCAGGTCGAC |
| Ubx2_S1 | GCAGCAGGTATTACGATAGAAGTATGTAATAGCTTTCATAGTGTAATC GAAGATGCGTACGCTGCAGGTCGAC |
| Ubx2_S2 | ACTCCAGAAACTCTTTGTACGCGTTTGTCGTTTTTAACGATATGCTATT TTATCAATCGATGAATTCGAGCTCG |
| Pdr5 S1 | AAAGACCCTTTTAAGTTTTCGTATCCGCTCGTTCGAAAGACTTTAGAC AAAAATGCGTACGCTGCAGGTCGAC |
| Pdr5 S2 | TAAAAAAGTCCATCTTGGTAAGTTTCTTTTCTTAACCAAATTCAAAATT CTATTAATCGATGAATTCGAGCTCG |

**Table S8** | Antisera used in this study.

| **Plasmid** | **Source** | **Identifier** |
|---|---|---|
| Rabbit polyclonal anti-Aco1 | *Ref. 69* | pTB104 |
| Rabbit polyclonal anti-Atp2 | *Ref. 70* | pTB104 |
| Rabbit polyclonal anti-Cdc48 | *Ref. 24* | GR5015 |
| Rabbit polyclonal anti-Cox5a | this study | GR1540 |
| Rabbit polyclonal anti-Hsp60 | *Ref. 69* | GR170 |
| Rabbit polyclonal anti-Ilv2 | this study | GR1009 |
| Rabbit polyclonal anti-Mdj1 | this study | GR1840 |
| Rabbit polyclonal anti-Msp1 | *Ref. 24* | GR1468 |
| Rabbit polyclonal anti-Om14 | *Ref. 24* | GR3040 |
| Rabbit polyclonal anti-Por1 | *Ref. 71* | GR3621 |
| Rabbit polyclonal anti-Pth2 | *Ref. 71* | GR797 |
| Rabbit polyclonal anti-Rip1 | *Ref. 69* | GR543 |
| Rabbit polyclonal anti-Rsp5 | *Ref. 41* | GR5063 |
| Rabbit polyclonal anti-Tim17 | this study | TB3015 |
| Rabbit polyclonal anti-Tom20 | *Ref. 72* | GR3225 |
| Rabbit polyclonal anti-Tom22 | *Ref. 72* | GR3227 |
| Rabbit polyclonal anti-Tom40 | *Ref. 72* | GR5104 |
| Rabbit polyclonal anti-Tom70 | *Ref. 72* | GR657 |
| Rabbit polyclonal anti-Ubp16 | *Ref. 71* | GR5020 |
| Rabbit polyclonal anti-Ubx2 | *Ref. 24* | GR1484 |
| Mmouse monoclonal anti-DHFR | Santa Cruz | sc-377091 |
| Mouse monoclonal anti-Dpm1 | Invitrogen | A-6429 |
| Mouse monoclonal anti-GFP | Santa Cruz | sc-9996 |
| Mouse monoclonal anti-HA | Santa Cruz | sc-7392 |
| Mouse monoclonal anti-Pgk1 | Santa Cruz | sc-130335 |
| Mouse monoclonal anti-ubiquitin | Santa Cruz | sc-8017 |
